# Seasonal and spatial variability of antibiotic resistance genes and Class I integrons in rivers of the Mekong Delta, Vietnam

**DOI:** 10.1101/2021.11.25.469999

**Authors:** Thi Thu Hang Pham, Khoa Dinh Hoang Dang, Emmanuelle Rohrbach, Florian Breider, Pierre Rossi

## Abstract

Aquaculture activities continue to expand in Vietnam, covering an estimated 700,000 ha, with 89% of these culture ponds located in the Mekong Delta. Since 2009, large-scale bacterial outbreaks have spread in response to this intensive farming. Antibiotics, even those considered a last resort, have only partially mitigated this problem. A side effect of the massive use of these chemicals is the appearance of mobile genetic elements associated with antibiotic resistance genes (ARGs). The large-scale emergence of a diverse bacterial resistome, accompanied by severe economic losses, has posed significant health risks to local residents. In this study, the seasonal and spatial distributions of the class I integrase (CL1) *intl*1 and the ARGs *sul*2 (sulfonamide), *BLA-oxa*1 (ß-lactams), and *erm*B (erythromycin) were quantified from water and sediment samples collected during two consecutive seasons along the Vam Co River and its tributary (Long An province, Vietnam). Results showed that CL1 was present in all river compartments, reaching 2.98×10^4^ copies/mL and 1.07×10^6^ copies/g of sediment, respectively. The highest relative copy abundances to the 16S rDNA gene were measured in water samples, with up to 3.02% for *BLA-oxa*1, followed by *sul*2 (1.16%) and *erm*B (0.46%). Strong seasonal (dry season vs. rainy season) and spatial trends were recorded for all resistance genes. Higher amounts of ARGs in river water could be associated with higher antibiotic use during the rainy season. In contrast, higher amounts of ARGs were recorded in river sediments during the dry season, making this habitat a potential reservoir of transient genes. Finally, the observations made in this study allowed us to clarify the environmental and anthropogenic influences that may promote the dispersal and persistence of ARGS in this riverine ecosystem.

**ABSTRACT ART:** 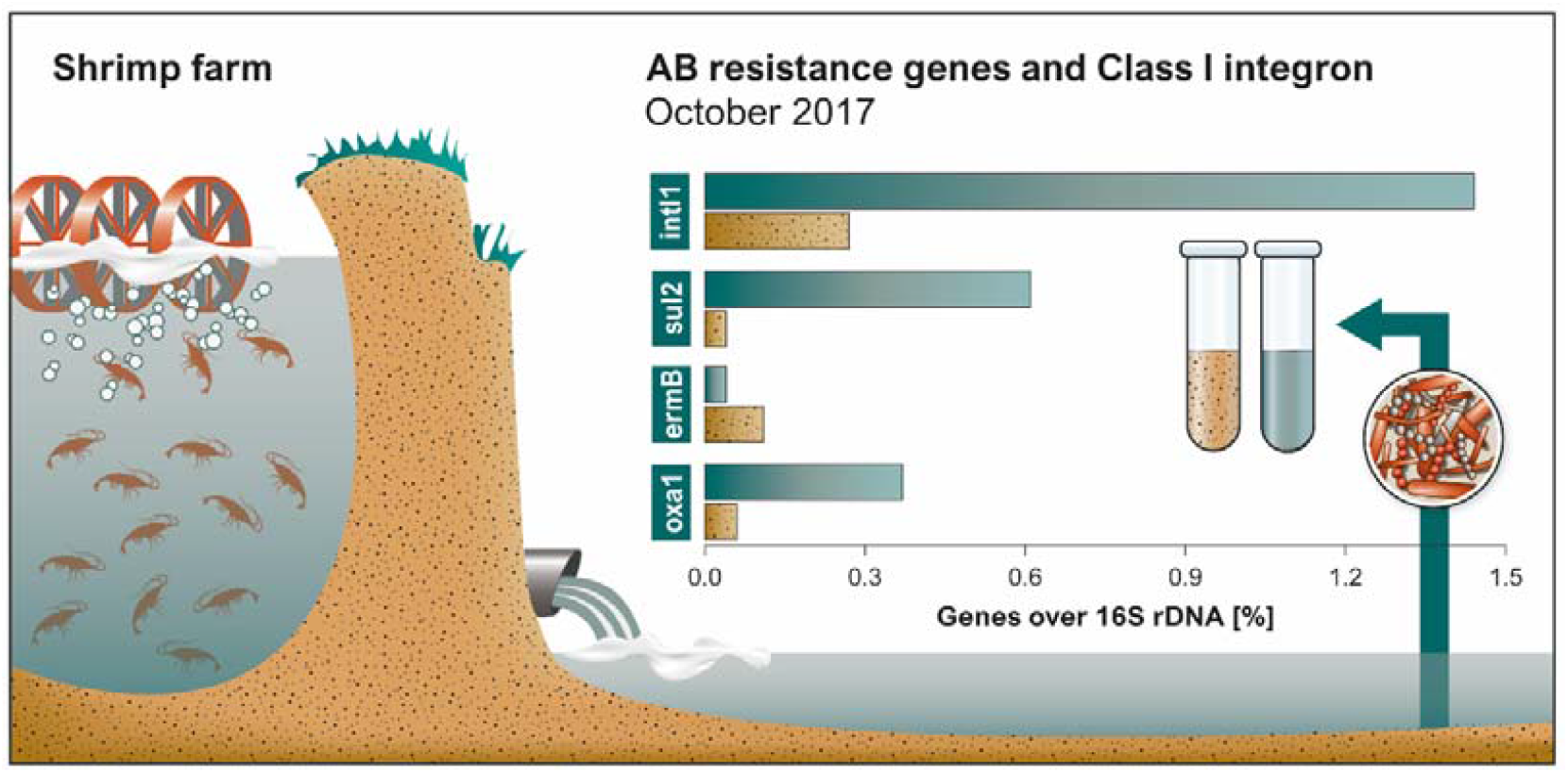

**HIGHLIGHTS:**

- In Vietnam, inland aquaculture massively relies on antibiotics to prevent epidemics
- Resistance genes were quantified along two rivers of the Mekong delta
- Seasonal (dry and rainy seasons) and spatial distributions were registered by qPCR
- BLA-*oxa1*and *sul2* reached highest abundances among bacterial communities
- Strong seasonal patterns and local variabilities were measured for CL1s and ARGs

## 1. INTRODUCTION

Aquaculture is considered one of the fastest growing food sectors and accounts for about 10% of all animal protein consumed worldwide (Thornber et al., 2020). According to recent FAO reports, production needs to increase by another 50% by 2050 while addressing the challenges of climate change and competition for natural resources (FAO, 2018). In 2000, the Vietnamese government implemented an agricultural diversification policy in the Mekong Delta, which shifted production from rice monoculture to a more diversified rice-based agricultural system by introducing aquaculture, horticulture, and fruit production. Aquaculture now plays an important role in the country’s economy, with shrimp exports peaking at $3.85 billion in 2017 (Anh et al., 2020). White shrimp (*Litopenaeus vannamei*) and black tiger shrimp (*Penaeus monodon*) are the dominant species cultivated in ponds covering an estimated 700,000 ha. About 89% of these ponds are located in the Mekong Delta (Hai et al., 2015, Anh et al., 2020). The success of shrimp farming has opened up opportunities for improved livelihoods, income, and employment for coastal and rural communities, providing approximately 4 million jobs nationwide (Lan, 2013; Khanh Nguyen et al., 2019; Kruse et al., 2020). More recently, and following the new Sustainable Development Goals, Viet Nam has defined an action plan to encourage farmers to reduce environmental impacts (Ngoc et al., 2021). This governmental decision follows the increasing environmental damage caused by this type of inland aquaculture (Pham et al., 2021). The implementation of these measures is also explained by the widespread cases of bacterial and viral diseases affecting the farming ponds.

Severe disease outbreaks have been the leading cause of loss in shrimp farming in recent decades (Duc, et al., 2015; Anh et al., 2020). Six viral diseases of crustaceans have the potential to cause substantial economic losses. Of these, the “White Spot Syndrome Virus” is the leading cause of high mortality rates in shrimp ponds and hatcheries (Millar et al., 2020; Thornber et al., 2020). Since 2009, substantial economic losses have been induced by the acute hepatopancreatic necrosis syndrome (AHPND) caused by *Vibrio parahaemolyticus*, a Gram-negative, halophilic, mesophilic, aerobic bacterium common in warm marine and estuarine habitats (Liu et al., 2019). Approximately 39,000 ha of shrimp ponds found in the Mekong Delta were affected by AHPND in 2011, with mortality rates up to 100%, inducing losses that were estimated to exceed $1 billion (FAO, 2013). Antibiotics (ABs), often applied directly to pond water, are the most common method to prevent and treat microbial outbreaks in shrimp farming and are currently an integral part of shrimp farming procedures (Chi et al., 2017; Binh et al., 2018; Ngoc et al., 2021). More than 30 antimicrobial compounds are approved in Viet Nam, including β-lactams, aminoglycosides, macrolides, tetracyclines, polymyxins, pleuromutilines, lincosamides, sulfonamides, and diaminopyrimidine (trimethoprim) (MARD, 2016, in Lulijwa et al., 2019). However, accurate data on antimicrobial sales and use are difficult to obtain because antibiotics are available over the counter and can be of variable quality (Tran et al., 2018). Due to limited access to diagnostic services, inadequate instructions for use, overuse, and poor education, treatment of an infection cannot be precisely targeted, with the risk of choosing an inappropriate or ineffective product (Lulijwa et al., 2019; Thornber et al., 2020). Over the past decade, the overuse of ABs has been identified as one of the major factors driving the increase in AB-resistant bacteria. The large-scale emergence of this bacterial resistome in aquatic habitats may not only result in severe economic losses, but also pose significant health risks to local communities. The spread of antibiotic resistance genes (ARGs) in potential and clinically relevant human and animal pathogens is hindered by local culture techniques and the use of ABs, even at low concentrations (Wistrand-Yuen et al., 2018). A growing body of literature describes the presence of ARGs in communities involved in aquaculture practices and the products themselves. For example, high levels of the *mcr*-1 gene conferring resistance to colistin in *Escherichia coli* have been detected within the microbiome of people living in aquaculture areas in China (Shen et al., 2018). Erythromycin resistance genes were recently detected by culture-independent techniques in commercial shrimp purchased from aquatic markets (Liu et al., 2019). In a previous study, we isolated and identified multidrug-resistant bacteria affiliated with the genera *Aeromonas, Acinetobacter, Citrobacter, Escherichia*, and *Klebsiella* from the Vam Co River (Long An Province, Viet Nam), a river receiving several channels of effluent from intensive shrimp farms (Pham et al., 2018). Sequencing of 41 plasmid-transmitted ARGs showed the presence of a wide range of ABs, including aminoglycosides, OXA-type ß-lactams, chloramphenicol, fluoroquinolone, macrolides, rifampicin, tetracycline, trimethoprim and sulfonamide. The association of ARGs with mobile genetic elements that enable their dissemination has been demonstrated worldwide (Gillings et al., 2008; Gao et al., 2012; Fang et al., 2021). Previous studies in the Mekong Delta have shown, for example, that *sul*2 genes have been detected in association with integrons and plasmids (Pham et al., 2018). Among all mobile genetic elements, Class 1 integrons are now considered key players in ARG dissemination, as they can easily acquire and express genes inserted into their structure as gene cassettes, expressed under a common promoter. These genetic elements are typically hosted on plasmids and transposons facilitating horizontal gene transfer (Preena et al., 2020). As shown by Gillings et al. (2015), Class 1 integrons provide a good measure of the overall level of antibiotic resistance genes. For example, recent studies have shown that the relative abundance of the *intl*1 gene correlates significantly with total AB concentrations in water and sediment (Chen et al., 2015).

However, and despite the potential for severe economic losses and significant health risks to local communities, very few studies have been conducted to assess the prevalence of the bacterial resistome in the Mekong Delta. The incidence of ARGs in microbial communities, their diversity, and their association with mobile elements such as integrons are all indications needed to understand the mechanisms of spread. In this project, we sought to understand how the quantity of selected ARGs fluctuates in natural streams affected by inland aquaculture practices. The novelty of this work is to observe in detail the seasonal and spatial evolution of these ARGs in relation to nearby human activities. Water and sediment samples were collected during two consecutive (dry and rainy) seasons along a tributary of the Vam Co River (Long An province, Viet Nam). *sul*2 (sulfonamide), *BLA-oxa1* (ß-lactams), *erm*B (erythromycin), and class I integrons were quantified by qPCR, normalized against 16S rDNA genes. Sampling sites included tributaries connected to small localities and effluent channels connected to shrimp farming ponds. The results obtained so far provide detailed information to accurately understand the prevalence and variability in time and space of the inland aquaculture-induced resistome in the Mekong Delta.

## 2. MATERIAL AND METHODS

### 2.1 Sampling

Two sampling campaigns were conducted in May and October 2017 on two rivers in the Mekong Delta, Vietnam. Sampling locations (SL) 1-8 were defined along an approximately 12 km transect on the Som Vam Co Tay River (Figure SI-1). Samples SL-9 and SL-10 were located on the Vam Co River, upstream and downstream of the junction with the aforementioned tributary, to assess the potential impact of mixing the two rivers. The section of the Som Vam Co Tay River that was selected for this study is bordered by large areas dedicated to shrimp farming. This section is particularly suitable for a detailed analysis of riparian microbial communities with potential impact from aquaculture activities. The SLs were selected based on the topology of the two rivers and were located near small tributaries and effluent outlets connected to intensive shrimp farming ponds. The Vam Co River, on the other hand, is bordered upstream by large areas dedicated mainly to agriculture (rice and dragon fruit), with the presence of two large industrial parks and a regional hospital (Table SI-1). For all SLs, one water sample and one sediment sample were collected within 5 meters of the riverbank (riverbank and sediment samples below). A second water sample was collected from the middle of the river (middle sample). Water samples were collected by hand from the water surface (approximately 20 cm deep) and stored in sterile 5-liter plastic containers. Sediment samples (approximately 0-5 cm deep) were collected manually with a bucket at a distance of approximately 10 meters from the river bank and transferred to sterile containers. All samples were shipped to the laboratory within 8 hours and stored at 4°C until processed. Dissolved oxygen, temperature, pH, electrical conductivity, and turbidity (expressed in nephelometric formazine units, FNU) were measured in the field using a YSI ProDSS portable multiparameter water quality meter (Xylem, USA) (Table SI-2 and SI-3). Note that sample SL-10 collected in the middle of the river (October) is missing.

### 2.2 Molecular analysis

DNA extraction from water and sediment samples was carried out as shown in Pham et al. (2018). Quantification of the ARGs, Class I integrase and 16S rDNA genes was carried out in triplicate in 10 μL reactions in a MIC qPCR Cycler (BioMolecular Systems, Australia) as follows: 2.5 mL template DNA, 2.1 μL water, 0.2 μL of each primer (100mM stock) (Table SI-4) and 5 μL of 2× KAPA SYBR Fast Universal qPCR kit (KAPA Biosystems, USA). Samples were cycled (40 cycles) at 95 °C for 10 s, followed by an extension at 62 °C for 30 s and acquisition at 72 °C for 20 s. The final melting step was carried out from 72 °C to 95 °C, at a rate of 0.1 °C/s. Analysis of the results was carried out using the built-in analytical software (micPCR, BioMolecular Systems, ver. 2.2). In average, efficiency (0.88 – 101.2%) and *r*^2^ values (> 0.995) were determined from eight points of the serial dilutions (10^8^–10^1^ copies) of each target gene. Based on calibration curves, Cq values were used to calculate the gene copy numbers which were normalized against mass (ng) of the extracted DNA.

## 3. RESULTS AND DISCUSSION

### 3.1 Physical and chemical measurements

Water samples collected from the Som Vam Co Tay tributary (SL-1 to SL-8) as well as those collected upstream and downstream of the junction with the Vam Co River (SL-9 and SL-10) were characterized by slightly acidic pH (pH 5.7 - 6.7) and oxygen depletion (OD 1.43 - 3.65 mg/L) (Table SI-2). No significant variation was noted between values measured in May and October, which were consistent with those measured locally in other Mekong tributaries (Strady et al., 2017). A seasonal difference was logically present in measured values for temperature, turbidity, and salinity due to heavy monsoon rains (Table SI-3). The mean temperature of the riverbank samples decreased from 31.4 to 29.2 °C in May and October, respectively. The values of the middle samples decreased from 31.3 to 29.9 °C during the same period. Turbidity values measured in May, averaging 18.6 and 19.5 FNU for the bank and middle samples, were significantly lower than those measured in October. During the wet season (October), higher flows increased the transport of solid particles. Turbidity values measured for the bank and middle samples averaged 145.8 and 62.4 FNU, respectively. It should be noted that the riverbank samples SL-3 and SL-5 contained large amounts of suspended particulate matter, which resulted in much higher turbidity records (542.6 and 380.0 FNU, respectively). These two samples also had much higher than average salinity, with 542 and 690 μScm^-1^ respectively (Table SI-2). Typically, during the rainy season, high salinity and turbidity values were recorded at the junction of the river with small streams and tributaries connected to the shrimp farming ponds. Intensive shrimp farming involves the rearing of 30-60 animals/m^2^, with maxima up to 200 animals/m^2^ (9 tons/ha) (Nguyen et al., 2019). Nutrient uptake is low under these conditions and a relatively small percentage (less than 20%) of nitrogen, phosphorus, and organic carbon is recovered from the biomass (Henares et al., 2019). To maintain water quality under conditions that allow for animal growth, common practices involve frequent pumping of 2-5% of the pond volume with removal of accumulated sludge waste from the bottom of the pond. This water is replaced by an identical volume of water accumulated in an adjacent settling tank, typically filled with relatively good quality river water pumped in at high tide. These volumes of effluent, rich in organic matter and salt, are discharged without further treatment to also compensate for heavy monsoon rains, in turn inducing eutrophication and salinization of the receiving surface waters (Lan, 2013). The bacterial cells in the culture ponds also enrich the natural microbial communities in the river sediments and surface waters (Shen, et al., 2020).

### 3.2 Quantitation of 16S rDNA genes

16S rDNA gene copies quantified in sediment samples showed significant differences between May and October (Figure 1). These numbers were significantly higher during the dry season (in May), ranging from 7.95×10^7^ to a maximum value of 2.14×10^8^ copies/g of sediment in SL-3 (Table SI-5). In October, copy numbers ranged from 1.77×10^7^ to 9.11×10^7^ copies/g. Only SL-1 showed nearly identical values in both seasons. For comparison, copy numbers measured in these sediment samples were close to those measured by Pham et al. (2018) in a previous dry season survey, with values ranging from 1.8×10^7^ to 2.1×10^8^ copies/g.

**Figure 1:**
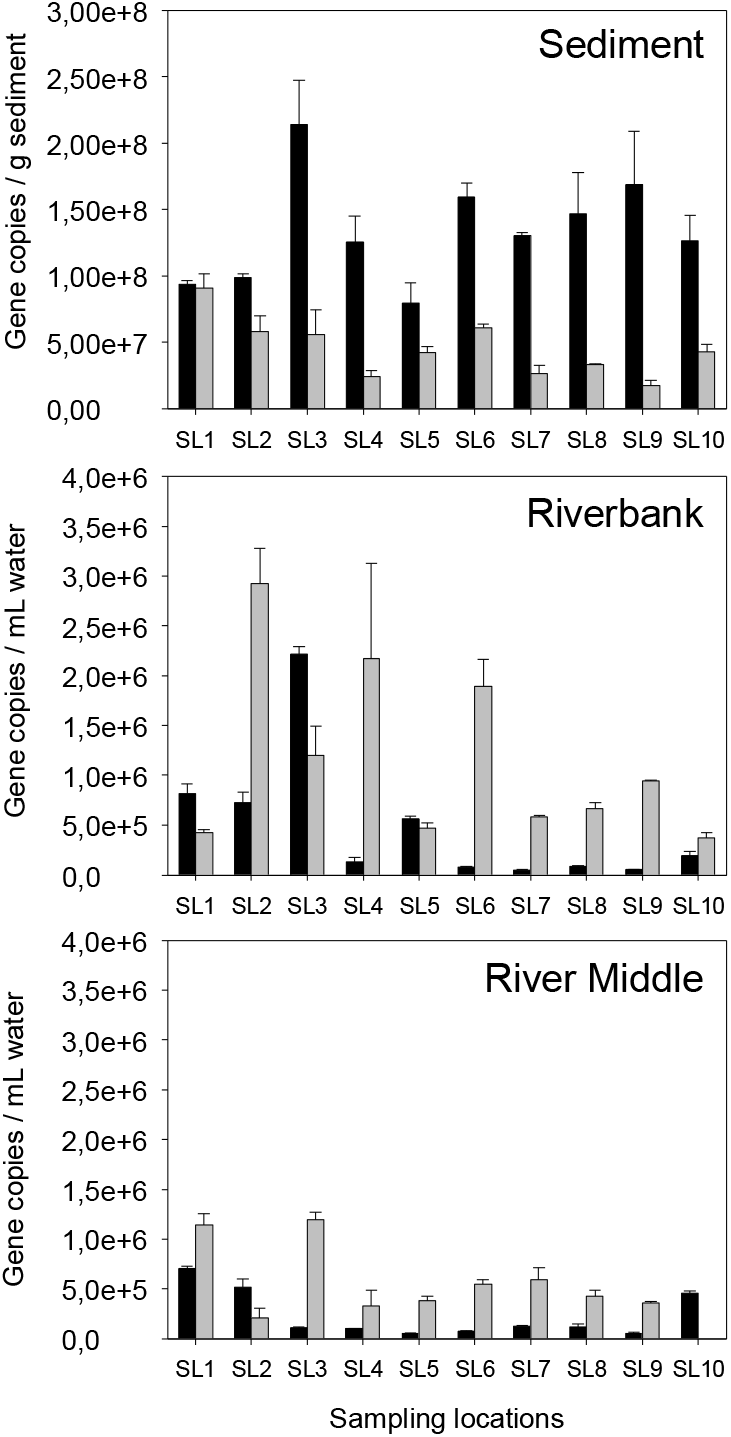
Copies of the 16S rDNA gene quantified in river sediment (top), riverbank (middle), and mid-river (bottom) samples. Black and gray bars: samples collected in May (dry season) and October (wet season), respectively.

Riverbank samples showed large seasonal differences in 16S rDNA gene copy number (Figure 1, middle). In October, variations between the lowest (SL-10: 3.71×10^5^ copies/g) and highest (SL-2: 2.92×10^6^ copies/g) copy numbers were spread over an order of magnitude. Conversely, the highest values (SL-2, SL-4, and SL-6) were located at the outlet of the shrimp tank effluent (Figure SI-1). These high values (>2.00×10^6^ copies/mL) may be related to culture practices, with the discharge of pond water to compensate for heavy monsoon rains. Classically, the highest copy numbers were recorded during the rainy season (October), with the exception of sites SL-1, SL-3 and SL-5 collected in May. Interestingly, all three of these sample sites were located near the outlet of small tributaries. Careful observation of the topography showed that these small rivers collected domestic effluent from small communities. In May, two orders of magnitude separated the highest value, SL-3 (2.22×10^6^ copies/mL) and the lowest, SL-7 (4.31×10^4^ copies/mL), indicating the presence of high spatial variability. For comparison, Pham et al. (2018) measured 2.1×10^5^ to 2.9×10^6^ copies/mL of the bacterial 16S rDNA gene for riverbank samples. These values are in agreement with those presented here. Results from samples collected in the middle of the river showed a very different pattern, including less seasonal variability (Figure 1, bottom). In May, the highest values were recorded for the two samples collected upstream, SL-1 and SL-2 with 7.05×10^5^ and 5.16×10^5^ copies/mL respectively, almost identical to those measured for the riverbank (Table SI-5). SL-5 showed the lowest value with 5.29×10^4^ copies/mL. During the wet season, values recorded were generally higher, ranging from 2.15×10^5^ (SL-2) to 1.19×10^6^ (SL-3) copies/mL. The low variability offered by the samples collected in the middle of the river is probably the result of constant mixing induced by the natural flow of the river, combined with tidal currents.

### 3.3 Quantitation of Class I integrons (CL1s)

The results of the quantitative analysis are shown in Figure 2 and the normalization ratios of *intI*1 copies to 16S rDNA are shown in Table 1. The complete quantitative data are provided in Table SI-6. The results show that *intI*1 was recovered consistently and frequently in large quantities in all river samples. In the sediment samples, the highest values were found among samples collected during the dry season, indicating a high seasonal pattern, reaching 1.07×10^6^ copies/mL in sample SL-9, an order of magnitude higher than the lowest amounts measured in sample SL-2 (Table SI-6). Sediment samples collected during the wet season showed less variation, with only about a threefold difference between samples SL-4 (3.70×10^4^ copies/mL) and SL-1 (1.98×10^5^ copies/mL). These values are comparable to those recorded in river sediments by studies conducted worldwide (Koczure et al., 2016; Sabri et al., 2020). However, the relative proportions of cells carrying this gene-capture system in the sediments were lower (0.30% and 0.27% on average for May and October, respectively) (Table 1) than those measured in other river systems (Amos et al., 2018).

**Figure 2:**
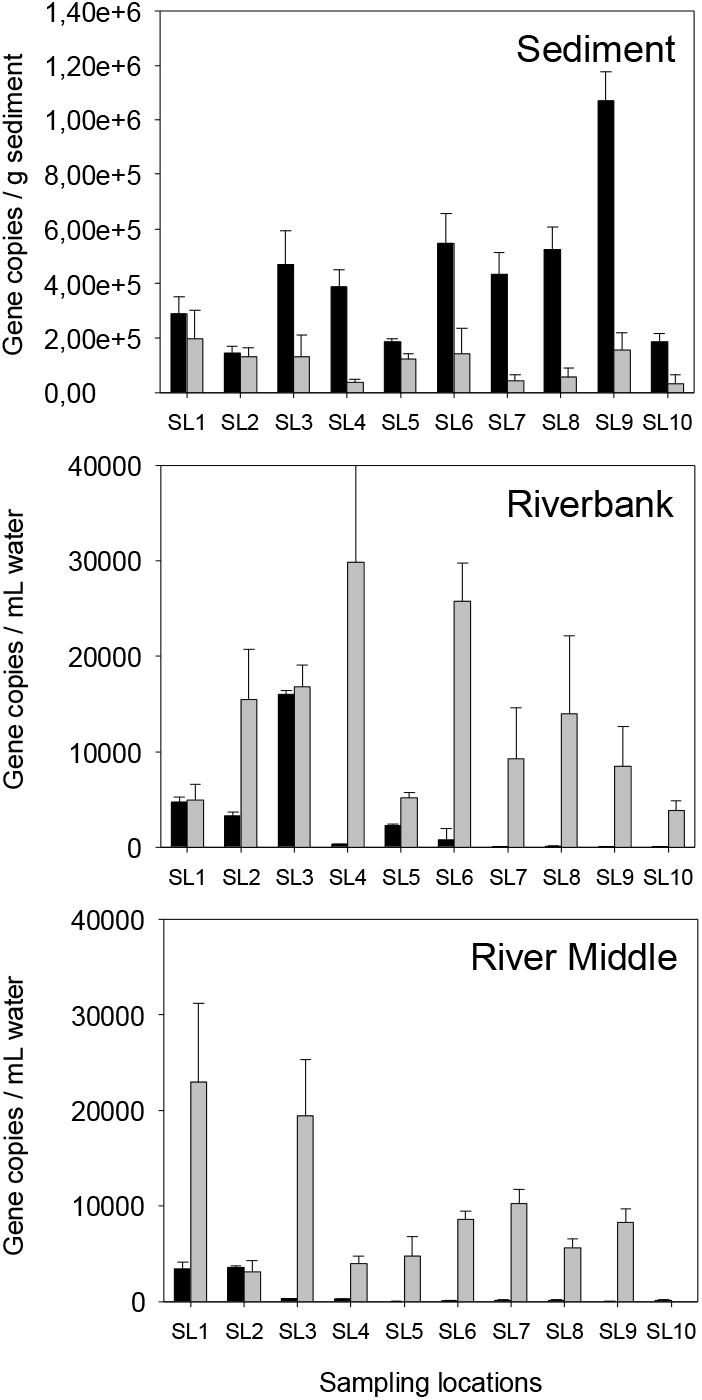
Copies of the *intI*1 gene quantified in river sediment (top), riverbank (middle), and mid-river (bottom) samples. Black and gray bars: samples collected in May (dry season) and October (wet season), respectively.

**Table 1:** Relative proportions of ARGs and Class 1 integron genes normalized to 16S rDNA gene copies.

| *intI1* | May 2017 |  |  | October 2017 |  |  |  | *sul2* | May 2017 |  |  | October 2017 |  |  |
| --- | --- | --- | --- | --- | --- | --- | --- | --- | --- | --- | --- | --- | --- | --- |
|  | Bank | Middle | Sediment | Edge | Middle | Sediment |  |  | Bank | Middle | Sediment | Edge | Middle | Sediment |
| SL-1 | 0.58% | 0.49% | 0.31% | 1.15% | 2.01% | 0.22% |  | SL-1 | 0.40% | 0.28% | 0.04% | 0.93% | 1.16% | 0.05% |
| SL-2 | 0.46% | 0.69% | 0.15% | 0.53% | 1.43% | 0.23% |  | SL-2 | 0.40% | 0.52% | 0.01% | 0.95% | 0.67% | 0.06% |
| SL-3 | 0.72% | 0.27% | 0.22% | 1.39% | 1.64% | 0.24% |  | SL-3 | 0.58% | 0.20% | 0.02% | 0.83% | 0.71% | 0.04% |
| SL-4 | 0.25% | 0.25% | 0.31% | 1.37% | 1.21% | 0.15% |  | SL-4 | 0.41% | 0.25% | 0.02% | 0.73% | 0.71% | 0.02% |
| SL-5 | 0.40% | 0.09% | 0.24% | 1.11% | 1.23% | 0.29% |  | SL-5 | 0.67% | 0.26% | 0.04% | 0.49% | 0.51% | 0.03% |
| SL-6 | 0.95% | 0.12% | 0.34% | 1.36% | 1.57% | 0.24% |  | SL-6 | 0.38% | 0.26% | 0.03% | 0.37% | 0.34% | 0.04% |
| SL-7 | 0.17% | 0.10% | 0.33% | 1.60% | 1.73% | 0.17% |  | SL-7 | 0.33% | 0.32% | 0.03% | 0.34% | 0.36% | 0.04% |
| SL-8 | 0.13% | 0.10% | 0.36% | 2.11% | 1.31% | 0.18% |  | SL-8 | 0.32% | 0.28% | 0.04% | 0.92% | 0.37% | 0.03% |
| SL-9 | 0.16% | 0.09% | 0.63% | 0.90% | 2.29% | 0.88% |  | SL-9 | 0.11% | 0.13% | 0.11% | 0.35% | 0.46% | 0.09% |
| SL-10 | 0.04% | 0.03% | 0.15% | 1.04% | ND | 0.08% |  | SL-10 | 0.22% | 0.13% | 0.03% | 0.26% | ND | 0.00% |
| MEAN | 0.38% | 0.22% | 0.30% | 1.26% | 1.60% | 0.27% |  | MEAN | 0.38% | 0.26% | 0.04% | 0.62% | 0.59% | 0.04% |

| *ermB* | May 2017 |  |  | October 2017 |  |  |  | *oxa1* | May 2017 |  |  | October 2017 |  |  |
| --- | --- | --- | --- | --- | --- | --- | --- | --- | --- | --- | --- | --- | --- | --- |
|  | Bank | Middle | Sediment | Edge | Middle | Sediment |  |  | Bank | Middle | Sediment | Edge | Middle | Sediment |
| SL-1 | 0.01% | 0.01% | 0.00% | 0.04% | 0.01% | 0.12% |  | SL-1 | 0.11% | 0.31% | 0.23% | 0.20% | 0.22% | 0.03% |
| SL-2 | 0.01% | 0.02% | 0.00% | 0.01% | 0.06% | 0.04% |  | SL-2 | 0.19% | 0.46% | 0.02% | 0.06% | 0.19% | 0.04% |
| SL-3 | 0.01% | 0.04% | 0.00% | 0.01% | 0.01% | 0.06% |  | SL-3 | 0.10% | 0.29% | 0.08% | 0.20% | 0.17% | 0.04% |
| SL-4 | 0.08% | 0.01% | 0.00% | 0.01% | 0.04% | 0.27% |  | SL-4 | 0.27% | 0.27% | 0.04% | 0.22% | 0.32% | 0.11% |
| SL-5 | 0.00% | 0.00% | 0.00% | 0.01% | 0.06% | 0.09% |  | SL-5 | 0.17% | 1.18% | ND | 0.20% | 0.62% | 0.08% |
| SL-6 | 0.00% | 0.01% | 0.00% | 0.02% | 0.07% | 0.06% |  | SL-6 | 1.84% | 2.72% | 0.03% | 0.10% | 0.45% | 0.06% |
| SL-7 | 0.46% | 0.20% | 0.00% | 0.06% | 0.12% | 0.06% |  | SL-7 | 3.02% | 1.38% | 0.06% | 0.87% | 0.53% | 0.07% |
| SL-8 | 0.10% | 0.00% | 0.00% | 0.02% | 0.01% | 0.13% |  | SL-8 | 1.57% | 1.24% | 0.04% | 0.01% | 0.87% | 0.03% |
| SL-9 | 0.00% | 0.03% | 0.00% | 0.01% | 0.04% | 0.19% |  | SL-9 | 1.13% | 0.94% | 0.10% | 0.56% | 0.73% | 0.06% |
| SL-10 | 0.00% | 0.05% | 0.00% | 0.07% | ND | 0.11% |  | SL-10 | 0.26% | 0.14% | 0.78% | 0.53% | ND | 0.04% |
| MEAN | 0.07% | 0.04% | 0.00% | 0.03% | 0.05% | 0.11% |  | MEAN | 0.86% | 0.89% | 0.15% | 0.29% | 0.45% | 0.06% |

The water samples showed strong seasonal fluctuations. Only a few samples, such as SL-3 with 1.60×10^4^ copies/mL (Table SI-6), had nearly identical copy numbers between the dry and rainy seasons (Table SI-6). All other samples collected in October (bank and middle) contained higher numbers of the CL1 gene. Differences between dry and wet seasons were remarkably high for samples Sl-4 through SL-10, with up to two orders of magnitude in gene copy numbers. The highest amounts were measured at the river’s bank in samples SL-4 and SL-6, with 2.98×10^4^ copies/mL and 2.58×10^4^ copies/mL, respectively. SL-1 and SL-3 showed the highest amounts for samples collected in the middle of the river, with 2.30×10^4^ copies/mL and 1.95×10^4^ copies/mL, respectively.

Interestingly, the relative proportions of *intI*1 gene copies (normalized to 16S rDNA gene copies) followed the same seasonal pattern, with an increase in all water samples from May to October (Table 1). This relative proportion increased on average from 0.38% to 1.26% for samples collected near the riverbanks, and from 0.22% to 1.60% for those collected in the middle of the river. SL-9 samples collected upstream of the junction with its tributary had the highest ratio (2.29%). The observed seasonal enrichment of bacteria carrying the CL1 gene capture system has been interpreted as the result of selective pressure on local communities. Recent studies have shown that the relative abundance of the *intI*1 gene correlated significantly with total AB concentrations in water and sediment (Chen et al., 2015). The seasonal increase observed in our study occurred concomitantly with the use of increasing amounts of ABs from August on in shrimp culture basins (Su et al., 2017; Pham et al., 2018). High rainfall events affect physical and chemical parameters of the pond water, such as temperature, salinity and pH, leading to deteriorating culture conditions. ABs are used in increasing quantities until the capture of mature animals, classically by the end of October. Nevertheless, the *intI*1 gene copies are three to four orders of magnitude lower than those found in Chinese static mariculture farms, a possible indication that river flow and tidal currents may play an important role in the constant washing of ABs and CL1-bearing bacterial cells to the ocean (Wang et al., 2019). The endemic presence of this gene-capture system is now documented in all environments disturbed by human activities. This is particularly the case when these habitats are impacted by the chronic presence of ABs, heavy metals and disinfectants (Amos et al., 2018; Oliveira-Pinto et al., 2018). These substances exert constant selection pressure on microbial communities, with concomitant rapid diffusion of various mobile elements carrying AB resistance genes across a wide range of bacterial species (Gillings et al., 2014). In our study, SL-9 (riverine environment) showed the highest relative proportion of the *intI*1 gene among riparian bacterial communities with 2.29%. Due to its geographical location, shrimp pond effluent may not be the only cause of the massive increase observed in this section of the Vam Co River. Other sources of anthropogenic activities, such as livestock and industrial effluents, municipal and hospital wastewater may also have contributed to the observed CL1 enrichment during the rainy season.

### 3.4 Quantitation of antibiotic resistance genes (ARGs)

Three genes responsible for antibiotic resistance were quantified in the sediment and water samples, namely *sul*2, *erm*B, and *BLA-oxa*1. The results of these analyses are presented in Figures 3, 4, and 5, and the values are provided in Tables SI-7 through SI-9. The relative proportions of normalized ARGs to 16S rDNA gene copies are presented in Table 1.

**Figure 3:**
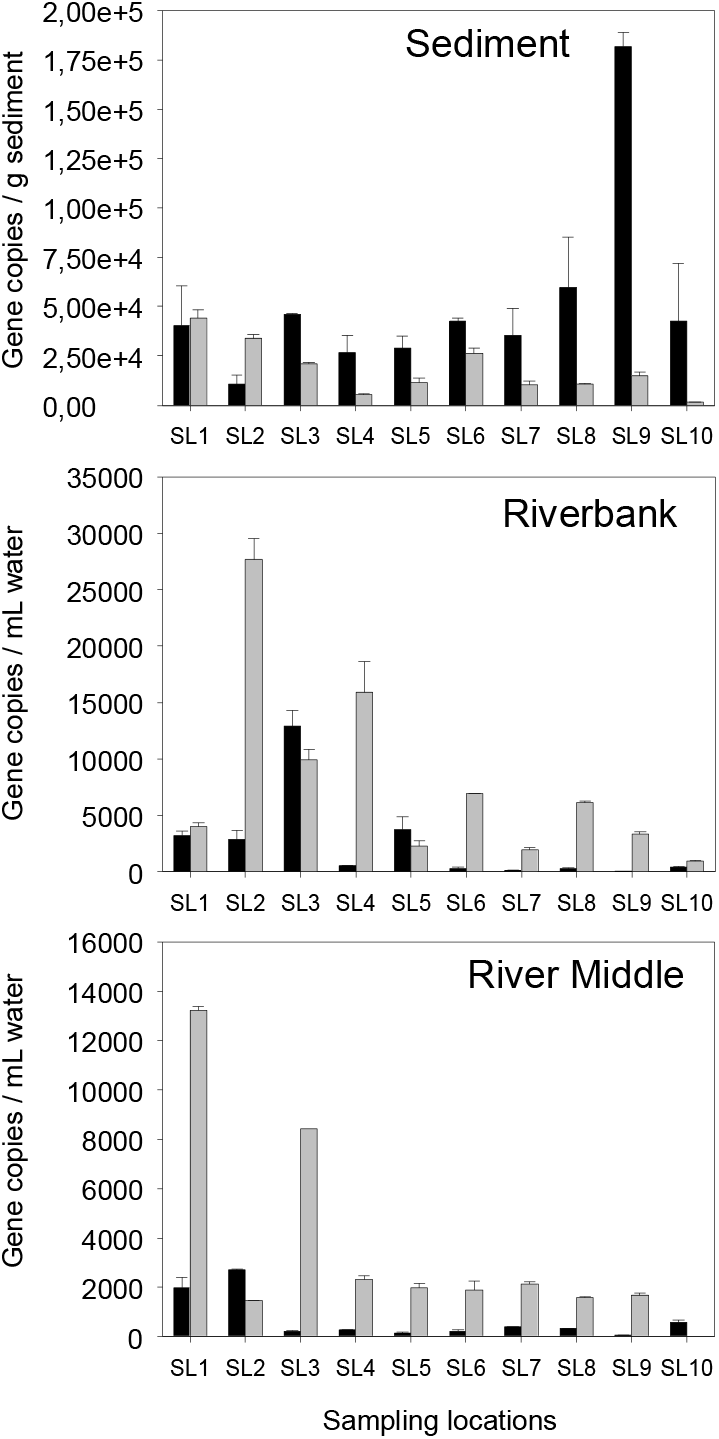
Copies of the *sul*2 gene quantified in river sediment (top), riverbank (middle), and mid-river (bottom) samples. Black and gray bars: samples collected in May (dry season) and October (wet season), respectively.

**Figure 4:**
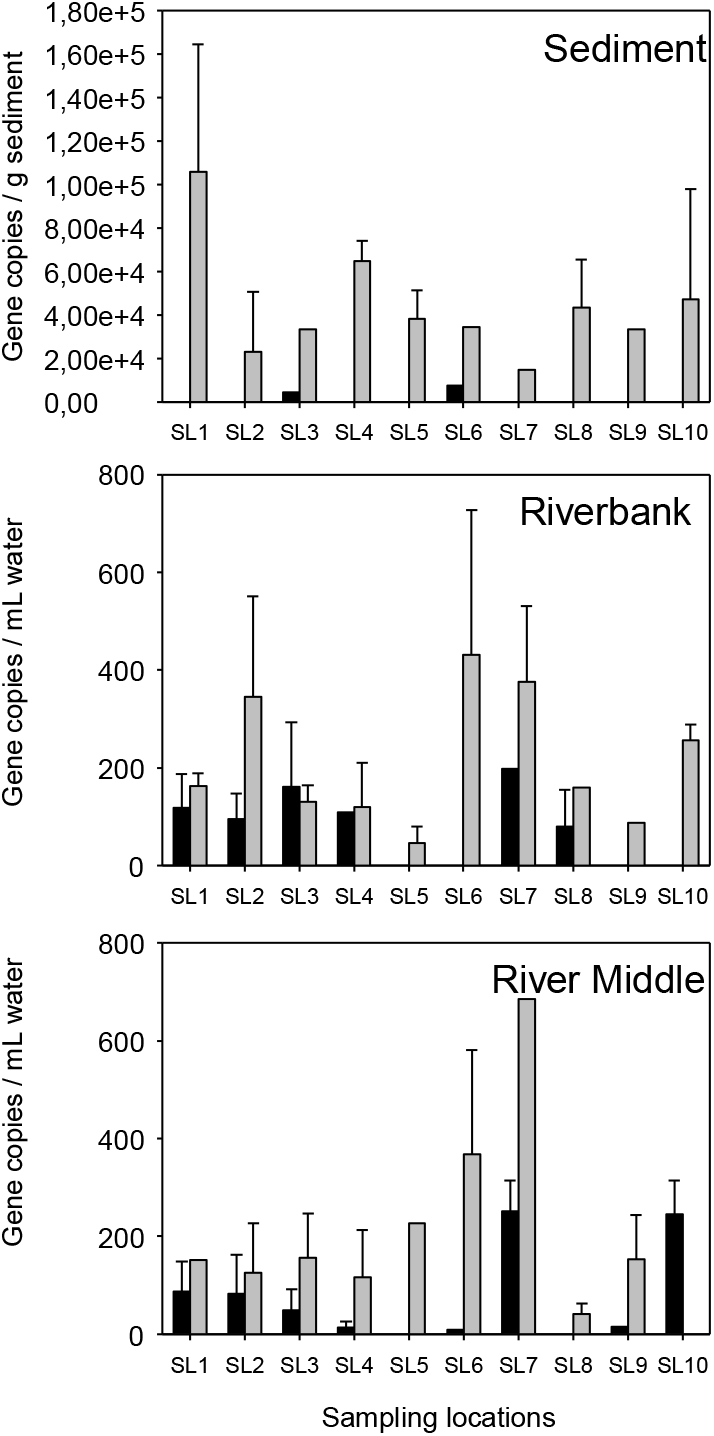
Copies of the *erm*B gene quantified in river sediment (top), riverbank (middle), and mid-river (bottom) samples. Black and gray bars: samples collected in May (dry season) and October (wet season), respectively.

**Figure 5:**
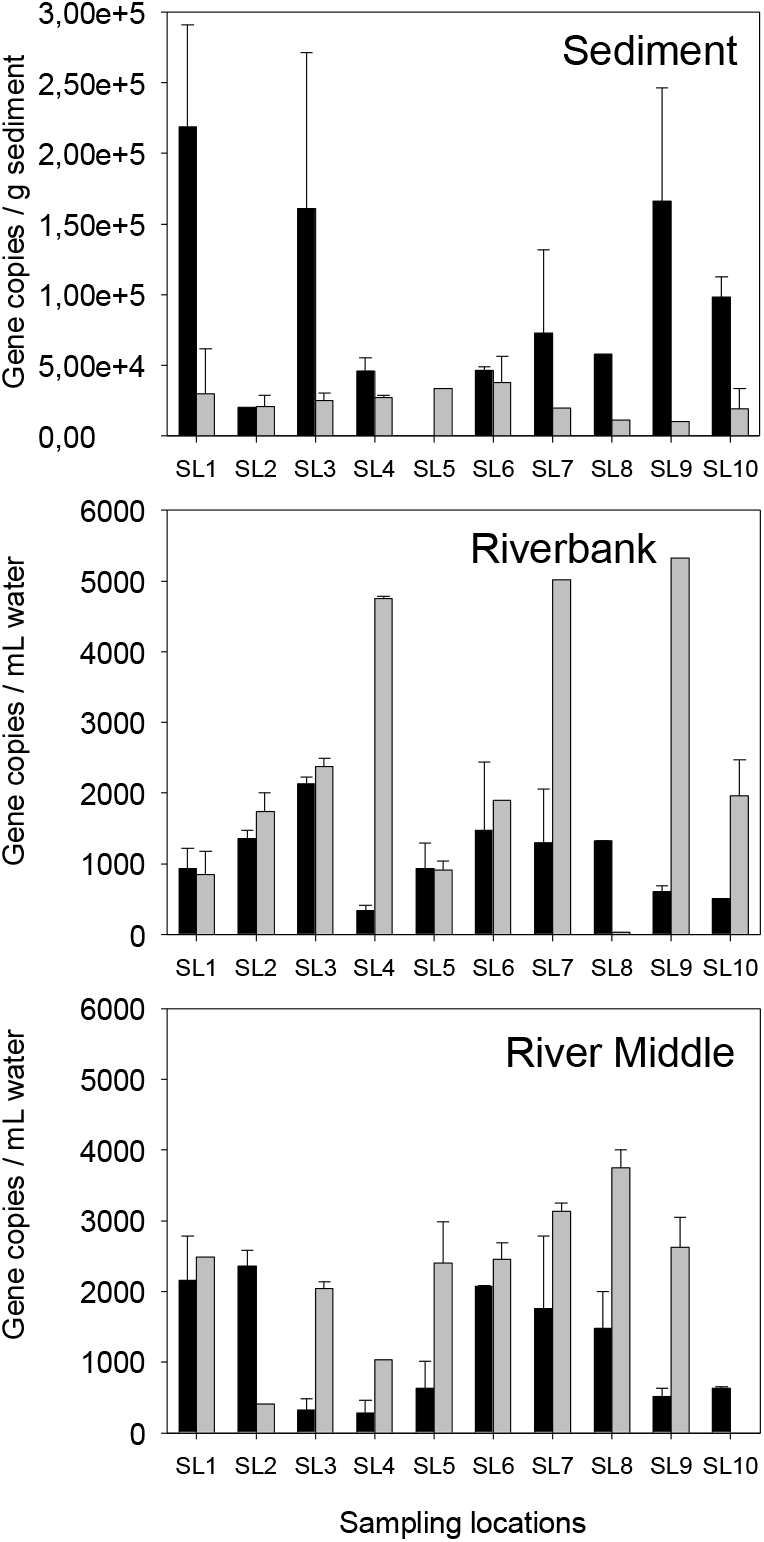
Copies of the BLA-*oxa*1 gene quantified in river sediment (top), riverbank (middle), and mid-river (bottom) samples. Black and gray bars: samples collected in May (dry season) and October (wet season), respectively.

Analysis of the results obtained for the *sul*2 gene showed spatial and seasonal patterns closely related to those observed for the *intI*1 gene (Table 1). Quantification of this resistance gene showed higher absolute numbers in sediment samples collected in May. Similarly, SL-9 showed the highest copy numbers, with 1.82×10^5^ copies/g. In October, copy numbers were generally lower, except for SL-1, with 4.41×10^4^ copies/g. However, and unlike CL1, the normalized relative proportions of *sul*2 showed no variation between the two seasons, with only 0.04%. This result demonstrated that this gene was uniformly maintained over the long term in the sediments of both river systems, albeit in small amounts.

Copies of the *sul*2 gene measured in water samples (riverbank) were highest in October, with significant local variability, with SL-2 reaching the highest copy number with 2.77×10^4^ copies/mL. SL-3 and SL-5 showed higher values only in May, with 1.29×10^4^ copies/mL and 3.78×10^3^ copies/mL, respectively. The mid-river water samples showed a similar seasonal pattern (Figure 3) with significantly higher copy numbers in October, particularly for SL-1 and SL-3, with 1.32×10^4^ and 8.43×10^4^ copies/mL, respectively. As shown for *intI*1, SL-2 showed the highest copy number in May with 2.70×10^3^ copies/mL. The relative proportions of *sul*2 gene copies in water samples, normalized to 16S rDNA gene copies, showed seasonal variation (Table 1). These proportions averaged 0.38% and 0.26% for bank and middle samples collected in May, respectively, and increased to 0.62% and 0.59% in October. Sample SL-1 (middle) had the highest relative proportion of *sul*2 at 1.16%. As a reminder, this sample concomitantly showed a significant increase in the proportion of *intI*1 genes, possibly indicating that a significant number of *sul*2 genes were carried by mobile genetic elements, as previously observed in rivers and estuaries (Sabri et al., 2020; Singh et al., 2019; Leng et al., 2020). In a study conducted in 2015, Chen et al. showed a significant correlation between *intI*1 gene abundance and dissolved AB concentrations, as well as with the abundance of *sul* genes, a possible consequence of their embedding in the same mobile genetic elements. The extensive use of sulfonamides in Viet Nam is confirmed by a 2013 study that showed that approximately 12 tons of sulfamethoxazole were discharged annually from the Mekong River into the South China Sea (Shimizu et al., 2013). These large quantities demonstrate the common use of sulfonamides for mitigation of bacterial diseases affecting aquaculture activities. In addition, it is now recognized that bacterial strains carrying a *sul* gene also develop resistance to other ABs, such as streptomycin, ampicillin, and trimethoprim, accentuating the emergence of multidrug-resistant strains.

In the present study, the use of this class of antibiotics resulted in the prevalence of resistances with a high degree of spatial heterogeneity. The results so far have confirmed that sulfonamid resistances are one of the major classes of ARGs in riverine habitats affected by shrimp farm effluents. However, the spread of *sul*2 in Mekong Delta rivers may also be correlated with other environmental factors. For example, Su et al. (2020) showed an excellent correlation between *sul*2 gene copies and COD measured in the farm ponds. Tide and river flow may increase the dilution rate of bacterial cells carrying this ARG, and at the same time spread these resistance genes throughout the river system. Nevertheless, the copy numbers measured in our study were much lower than those measured in other intensive aquaculture systems. Wang et al. (2019) measured 3.5×10^7^ to 6.5×10^10^ copies/mL of the *sul*2 gene in a Chinese salmon mariculture system, values that exceed the copy numbers measured in our study by more than 4 orders of magnitude.

Erythromycin is a macrolide listed as a compound of critical importance to human health (WHO, 2017). A corresponding resistance gene, *erm*B, was frequently detected in multidrug-resistant bacterial strains characterized in a previous campaign conducted on an identical section of the Vam Co River (Pham et al., 2018). Interestingly, the prevalence and distribution patterns of this ARG in the sediment samples differed significantly from the results obtained for *intI*1 and *sul*2 (Figure 4). The prevalence was reversed, with significantly higher copy numbers in the wet season (October) than in the dry season, with a maximum of 1.06×10^5^ copies/g in sample SL1 (Table SI-8). During the dry season (May), only SL-3 and SL-6 showed quantifiable values, with 4.31×10^3^ and 7.33×10^3^ copies/g, respectively. The relative proportions of 16S rDNA genes were highest in October among the microbial communities populating the river sediments, with a maximum of 0.27% for SL-4 (Table 1). The seasonal trend was also strong among water samples, but with significantly lower copy numbers than those measured for *intI*1 and *sul*2. SL-6 and SL-7 had the highest copy numbers for both bank and middle samples, with values ranging from 368 to 6845 copies/mL (Table SI-8). Like many other *erm* genes, *erm*B, is considered one of the most prevalent resistance genes in environmental microorganisms (Pham et al., 2018). Recently, *erm*B was detected at approx. 10^4^ copies/gr of sediment, and 10^2^-10^3^ copies/ml in wastewater samples (Sabri et al., 2020). This erythromycin resistance gene was found among the bacterial microflora populating the digestive tract of *Penaeus vannamei* specimens purchased from markets (Liu et al., 2019). In the present study, the strong seasonal trend shown by this ARG may indicate that the use of this class of ABs was limited to the rainy season, when the risk of bacterial infections is highest. Interestingly, only two sediment samples collected in May had quantifiable amounts of *erm*B genes. The reasons why the prevalence of this gene was reduced in the May-collected sediment samples could not be inferred from our data set. Nevertheless, previous studies have shown that this ARG can be easily removed from a microbial community, as is the case during biological treatment (Di Cesare et al., 2016).

The BLA*-oxa*1 enzymes compose one of the major plasmid-mediated β-lactamases involved in amoxicillin-clavulanic acid resistance. Among these genes, class D carbapenemases consist of OXA-type β-lactamases, the most versatile family of β-lactamases (Pham et al., 2018). As a reminder, carbapenems are currently used as last-line ABs to treat resistant bacterial infections and are threatened by β-lactamases (WHO, 2017). Our results showed that copy numbers of this class of ABs, considered “antibiotics of last resort,” were found in all river compartments. In the sediment samples, the results showed a marked seasonal pattern closely related to those observed for *sul*2 and *intI*1 (Figure 5). Copy numbers measured during the dry season were generally the highest, with a maximum of 2.19×10^5^ copies/g in SL-1. It should be noted that, as with *sul*2 and *intI*1, high amounts were also measured in the Vam Co River samples, with 1.66×10^5^ copies/g in SL-9. With the exception of SL-1 (0.23%), the relative proportions calculated from the 16S RNA genes showed little local and seasonal variation, averaging 0.08% and 0.06% for May and October, respectively (Table 1). These numbers indicate that the river sediments are a stable reservoir for this RNA, even though it has a low prevalence among bacterial communities.

BLA-*oxa*1 copy numbers measured in water samples were highest during the rainy season and in samples collected near the bank. Samples SL-4, SL-7, and SL-9 were distinguished by amounts near 5×10^3^ copies/mL. Samples collected in the middle of the river had a more homogeneous pattern, with lower copy numbers, ranging from 4.12×10^2^ (SL-2) to 3.74×10^3^ (SL-8). In terms of copy number, this ARG showed a seasonal fluctuation of low amplitude, similar to that demonstrated by *erm*B. The relative proportions of BLA-*oxa*1 to 16S rDNA genes revealed a seasonal pattern, with higher proportions in May than in October (Table 1). Sample SL-7 had a significant 3.02% of this ARG in the river water system. The persistent presence of the broad-spectrum BLA-*oxa*1 resistance genes, albeit in low amounts in Mekong Delta surface waters, is of great concern. The higher average prevalence of this gene during the dry season cannot be explained solely by hypothetical illegal use of carbapenems in aquaculture activities, as these compounds have never been approved for use in farming in any country in the world (Bonardi & Pitino, 2019). The prevalence in SL-9 could reflect the use of this class of ABs in other human activities, for example in hospitals located upstream of the geographical area of interest (Hoa et al., 2019). This constant presence induces a memory effect in these natural compartments, which is likely to preserve and eventually transmit resistance throughout the ecosystem. The most likely explanation for this memory effect is co-occurrence with other ARGs, as well as metal and disinfection resistance genes (Di Cesare et al., 2016). Uncontrolled use of other antibiotics possibly induces selection pressure favorable to the preservation of mobile genetic elements, jointly carrying the genes responsible for carbapenem resistance (Hamza et al., 2020).

## 4. CONCLUSIONS

Aquaculture-related activities continue to expand in Vietnam, and the outlook is favorable for shrimp production (Khanh Nguyen et al., 2019). However, the massive increase in area dedicated to shrimp monoculture also increases the risk of large-scale bacterial outbreaks. The massive use of ABs, even those considered a possible last resort, only partially mitigates this problem. Side effects of the use of these products include the appearance of large quantities of mobile genetic elements associated with resistance genes among the bacterial communities populating the culture ponds.

There is a growing awareness that these genetic elements, such as CL1s, contribute to the spread of ARGs in ecosystems adjacent to those affected by agricultural operations. In the present study, large spatial and seasonal variations were observed for ARGs and Class I integrons in the Mekong Delta river system. The results showed that the apparent distribution of ARGs and CL1s in the river system was not uniform. The prevalence of the quantified genes differed both in habitat (water vs. sediment) and in time (dry vs. wet season). The distribution of small tributaries and outfalls from the growing basins could, logically, explain the spatial variability in ARGs, and particularly that observed in the riverbank samples. A more precise location of the growing basins and their tributaries, as well as any other human activity involving the use of RGW, would certainly refine the understanding of the system. In addition, seasonal variability results from the action of environmental parameters that will require more attention in the future. These parameters are likely to favor both dispersal and persistence of the resistome in Mekong Delta tributaries. Strong currents and tides may have played a key role in the observed seasonal decrease for resistance genes that emerged during the rainy season. Other environmental variables, such as high organic matter loads, may also have played a role (Su et al., 2020). In addition to these environmental factors, there are logically human factors, as shrimp farming is a labor-intensive activity, resulting in a high prevalence of multidrug-resistant strains among workers. Finally, the persistent presence of ARGs and CL1s along the Vam Co River and its tributary confirmed that water and sediment are routes of dissemination, highlighting the urgent development of alternative agricultural practices.

## 5. ACKNOWLEDGEMENTS

This research was funded partly by the Cooperation and Development Center (CODEV) at EPFL, and by the Vietnam National University Ho Chi Minh City (VNU-HCM). We acknowledge Ho To Thi Kha Mui, Tran Phuong Anh and Pham Ngoc Tu Anh from the VNU-HCM for their technical assistance.

## 6. DATA ACCESSIBILITY

Full data and metadata sets are available on Zenodo: https://doi.org/10.5281/zenodo.5727167

