## Supplementary_Information for "Seasonal and spatial variability of antibiotic resistance genes and Class I integrons in rivers of the Mekong Delta, Vietnam"

### A. **Sampling locations**

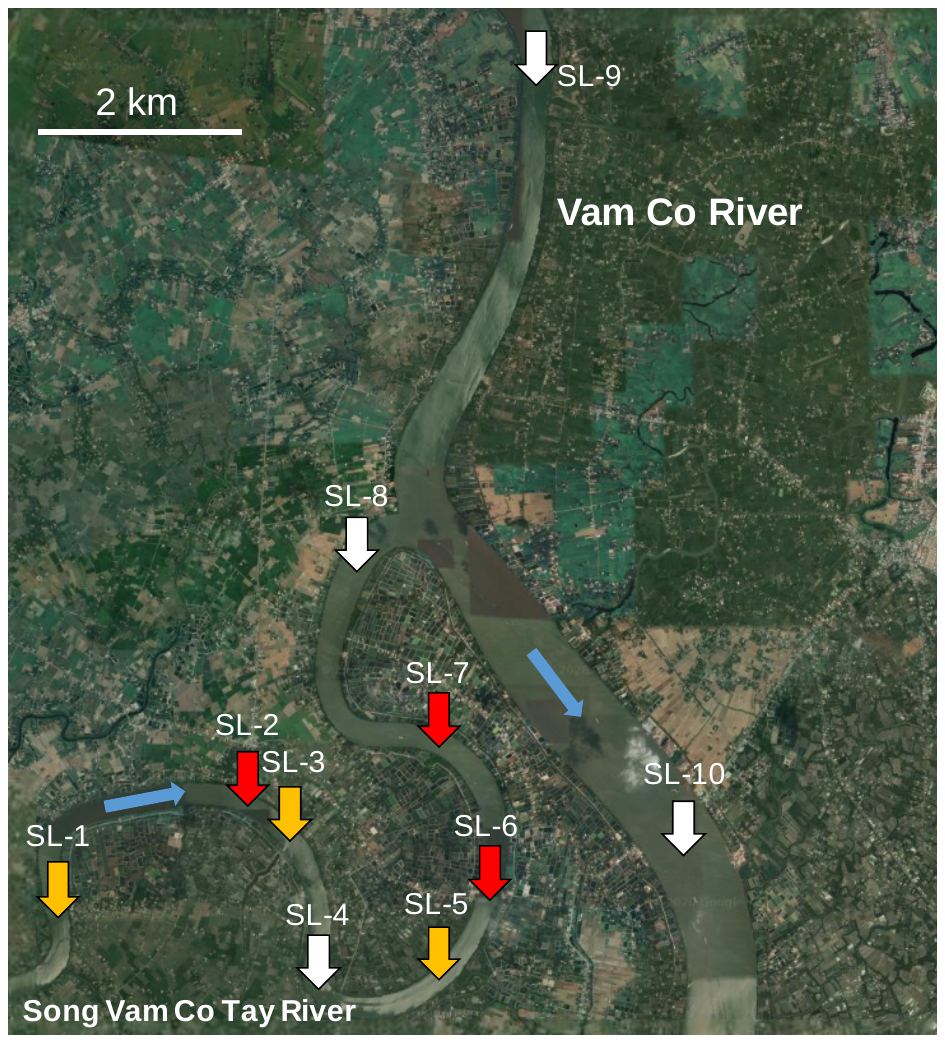

**Figure SI-1**: Geographic location of the ten sampling points (SL) collected on the Vam Co River and its tributary (Long An province, Vietnam). Two water samples (riverbank and mid-river) and one sediment sample were collected at each location during two sampling campaigns (May and October 2017). Red arrows correspond to SLs in close proximity to the shrimp pond effluents. Orange arrows indicate SLs located at the outlets of small tributaries. Stream directions are indicated by a blue arrow. Image from Google Maps.

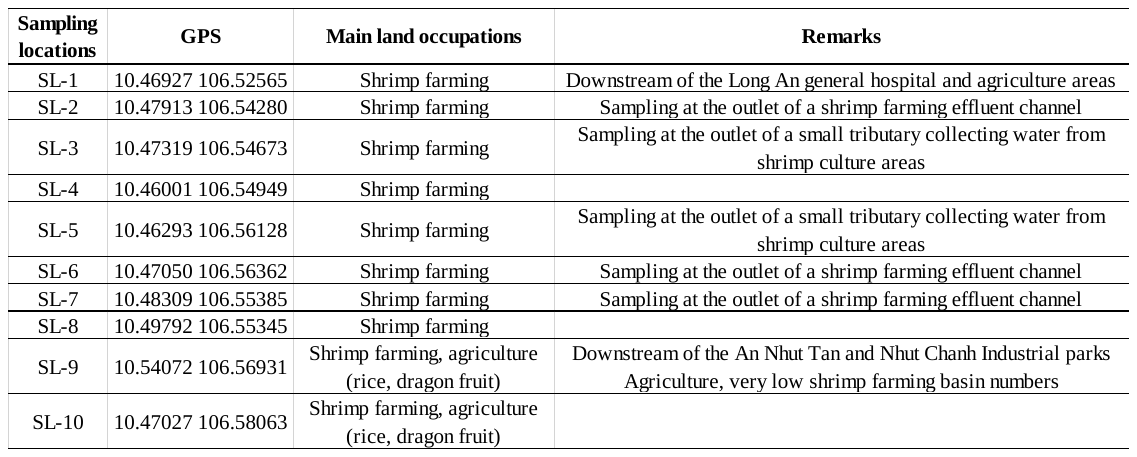

**Table SI-1**: Sampling locations with GPS coordinates (Decimal Degrees) and main land occupations.

### B. **Physical and chemical data set**

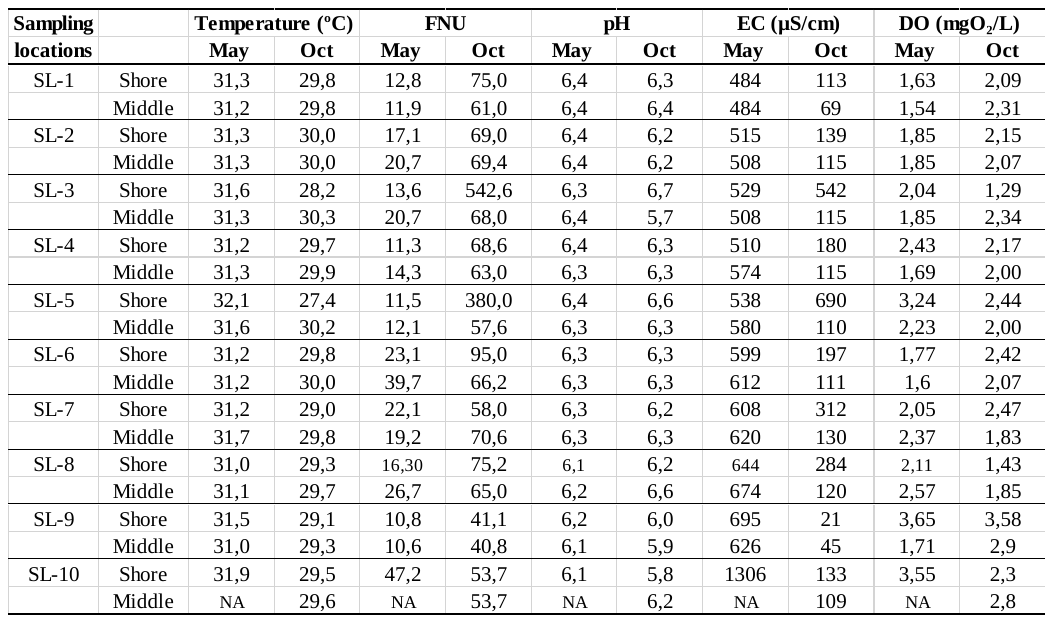

**Table SI-2:** Physical and chemical parameters measured for water samples. Shore and Middle: samples respectively from the shore and from the middle of the river. FNU: Formazine Nephelometric Unit.

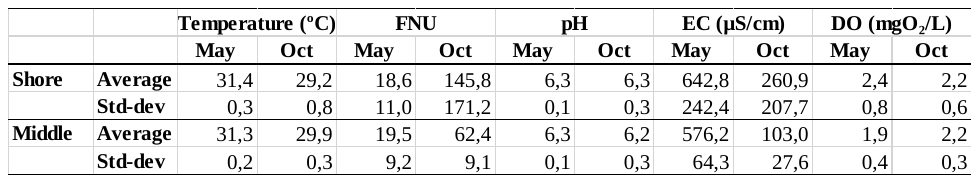

**Table SI-3:** Average and standard deviation computed from Table SI-2. FNU: Formazine Nephelometric Unit.

### C. **Primer list**

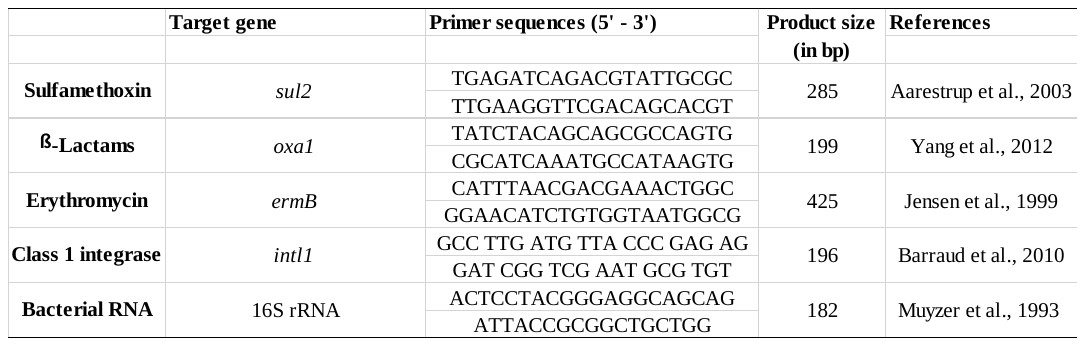

**Table SI-4:** Primers used in quantitative PCR.

### D. **Data sets obtained from quantitative assessments**

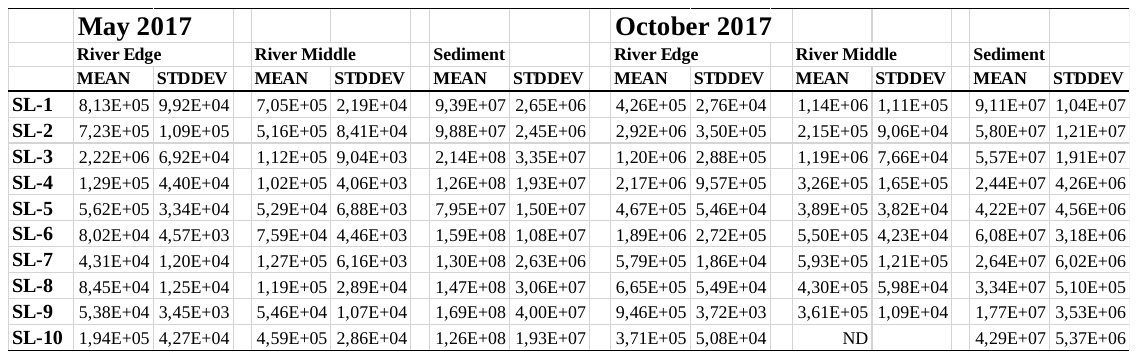

**Table SI-5:** 16S rDNA gene copy number measured in river water samples (edge and middle) and river sediment. ND: not determined.

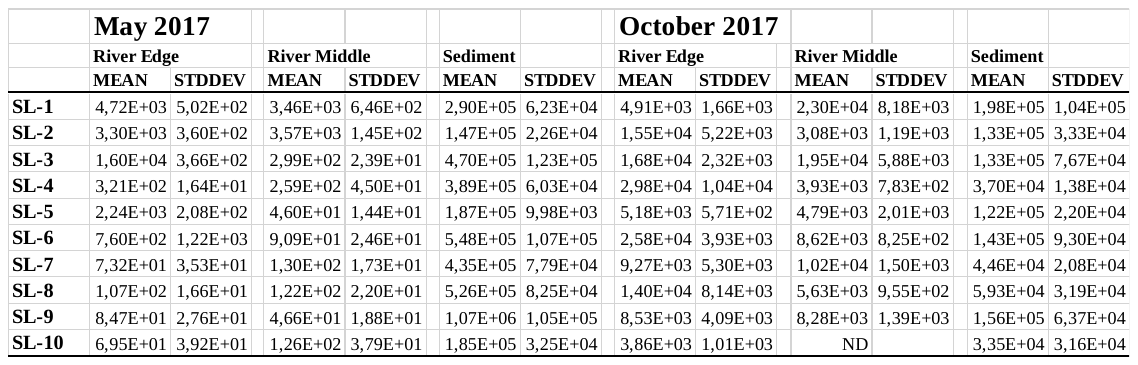

**Table SI-6:** *intl*1 gene copy number measured in river water samples (edge and middle) and river sediment. ND: not determined..

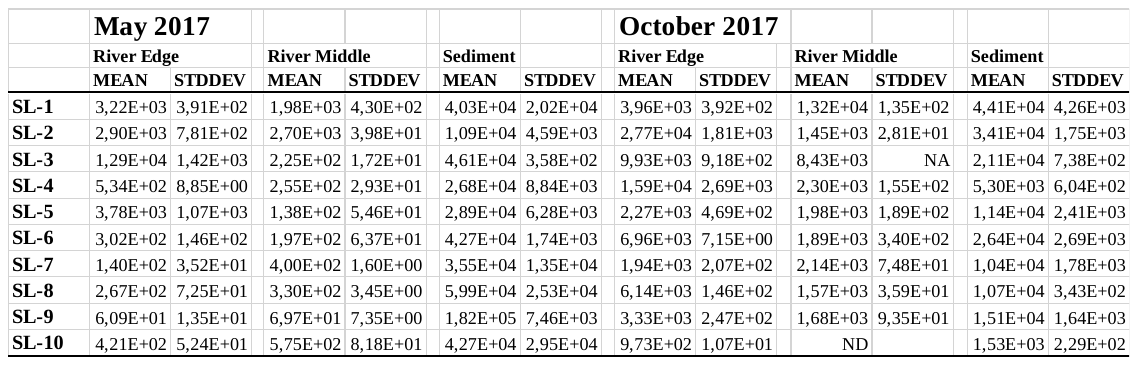

**Table SI-7:** *sul*2 gene copy number measured in river water samples (edge and middle) and river sediment. ND: not determined. NA: missing replicate.

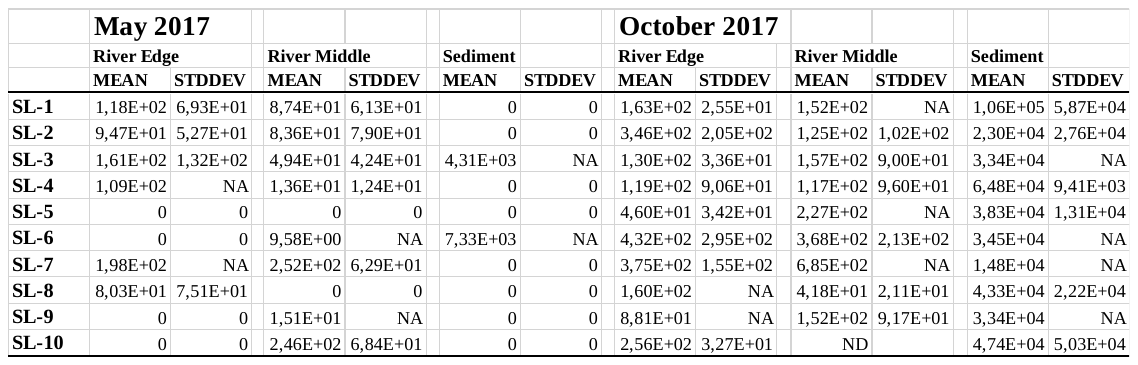

**Table SI-8:** *erm*B gene copy number measured in river water samples (edge and middle) and river sediment. ND: not determined. NA: missing replicate.

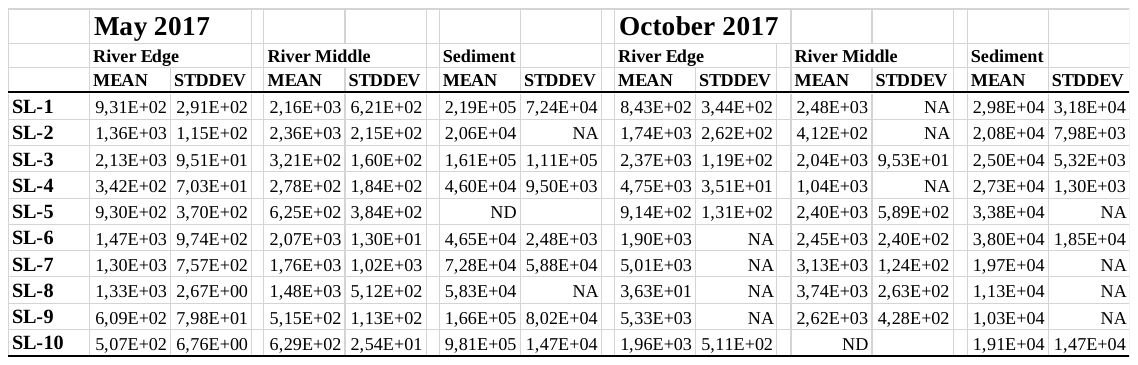

**Table SI-9:** *oxa*1 gene copy number measured in river water samples (edge and middle) and river sediment. ND: not determined. NA: missing replicate.

### E. **Relative percentages**

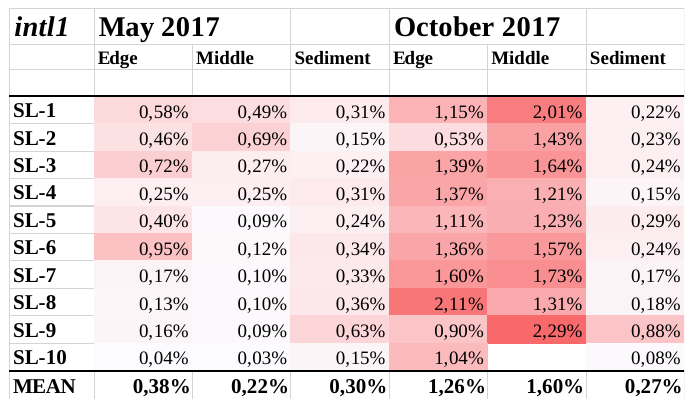

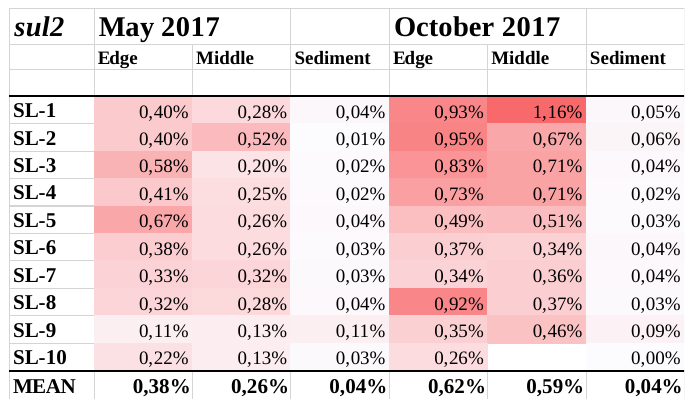

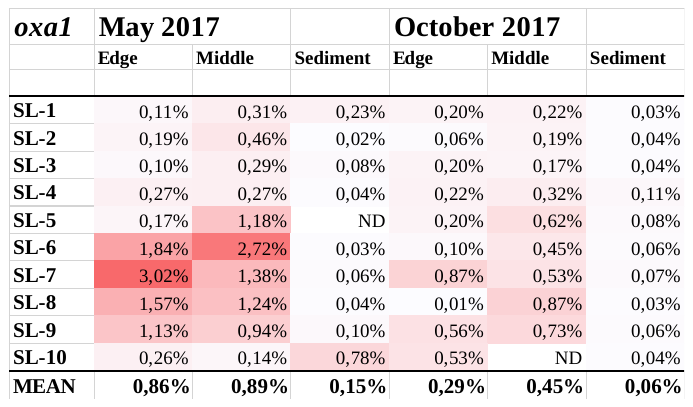

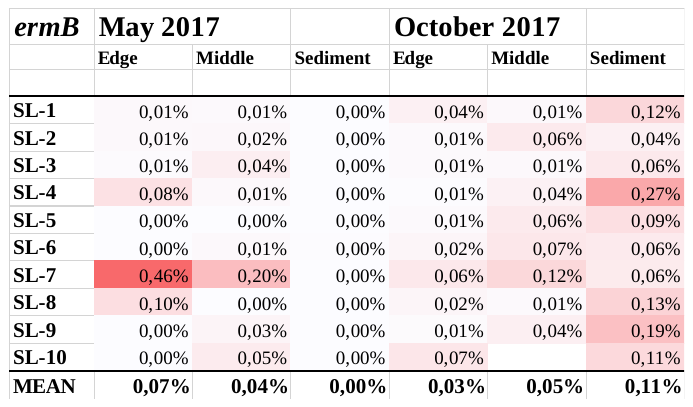

**Table SI-10:** Relative proportions computed for *intl1*, *sul2*, *ermB* and *oxa*1 using gene copy numbers normalized to 16S rDNA gene copies.

### F. **References**

Aarestrup, F.M., Lertworapreecha, M., Evans, M.C., Bangtrakulnonth, A., Chalermchaikit, T., Hendriksen, R.S., & Wegener, H.C. 2003. Antimicrobial susceptibility and occurrence of resistance genes among *Salmonella enterica* serovar Weltevreden from different countries. *Journal of Antimicrobial Chemotherapy*, *52*(4), 715-718.

Barraud, O., Baclet, M.C., Denis, F., Ploy, M.C. 2010. Quantitative multiplex real-time PCR for detecting class 1, 2 and 3 integrons. *Journal of antimicrobial chemotherapy*, 65(8), 1642-1645.

Jensen, L. B., Frimodt-Møller, N., Aarestrup, F. M. 1999. Presence of erm gene classes in gram-positive bacteria of animal and human origin in Denmark. *FEMS Microbiology Letters*, *170*(1), 151-158.

Muyzer G., Dewaal E.C., Uitterlinden A.G. 1993. Profiling of complex microbial populations by denaturing gradient gel electrophoresis analysis of polymerase chain reaction amplified-genes coding for 16S rRNA. *Appl. Environ. Microbiol*. 59(3): 695-700.

Yang, Y., Zhang, T., Zhang, X.X., Liang, D.W., Zhang, M., Gao, D.W., … & Fang, H.H. 2012. Quantification and characterization of β-lactam resistance genes in 15 sewage treatment plants from East Asia and North America. *Applied microbiology and biotechnology*, 95(5), 1351-1358.
